# Mechanically competitive regulation of cell volume in cytoplasm-sharing cells connected by intercellular bridges

**DOI:** 10.64898/2026.01.26.701669

**Authors:** Hiroshi Koyama, Kanako Ikami, Lei Lei, Toshihiko Fujimori

**Affiliations:** Division of Embryology, National Institute for Basic Biology, 5-1 Higashiyama, Myodaiji, Okazaki, Aichi 444-8787, Japan; SOKENDAI (The Graduate University for Advanced Studies), 5-1 Higashiyama, Myodaiji, Okazaki, Aichi 444-8787, Japan; Howard Hughes Medical Institute, University of California Davis, Davis, California, CA 95616; Department of Microbiology & Molecular Genetics, University of California Davis, Davis, CA 95616; Department of Obstetrics, Gynecology and Women’s Health, University of Missouri School of Medicine, Columbia, MO 65211; Division of Biological Sciences, College of Arts and Sciences, University of Missouri, Columbia, MO 65211

**Keywords:** mouse germline cyst, Laplace pressure, cell-cell contact, cytoplasmic flow, mathematical modeling

## Abstract

In multicellular organisms, various cellular structures exhibit cytoplasmic sharing, where cells remain interconnected. While essential for development and function in contexts such as germ cell formation and insect early embryos, the physical basis of cell volume regulation in these systems remains poorly defined. Germline cysts are formed by interconnected sister cells via intercellular bridges. In mice, germline cysts form during gametogenesis in fetal ovaries and testes. In mouse fetal female cysts, cells with numerous bridges preferentially differentiate into oocytes by selectively increasing their volume, a process that may be mediated through cytoplasmic flow. This volume bias may be influenced by hydrostatic pressure within the cytoplasm. Here, we theoretically investigate how the mechanical properties of cells affect cytoplasmic pressure and volume distribution within interconnected cells. Our soap-bubble model revealed that cells with more bridges exhibit increased volume when they have large cell-cell contact areas, as observed in fetal cysts. We found that incorporating cell cycle (including cell growth and cell division) significantly enhances the likelihood of volume bias in favor of cells with more bridges. These theoretical findings suggest that intrinsic mechanical properties, coupled with cell cycle, establish robust cyst development in fetal female germline cysts. Our findings also provide insights into the volume dynamics observed in adult male germline cysts, which are characterized by smaller cell-cell contact areas.

**Impact statement:** A theoretical model demonstrates how mechanical properties and cell cycle dynamics regulate volume distribution in germline cysts, providing a physical basis for oocyte differentiation.

## Introduction

In multicellular organisms, cytoplasm in each cell is typically separated by cell membranes. On the other hand, there are some cell types whose cytoplasm is shared among multiple cells. Notable examples include the early syncytial embryos in insects like *Drosophila* during cellularization where the nuclei are partially separated by cell membrane (Lv et al., 2021); the formation of oocytes from the gonad in nematodes where each germ cell is connected to a central cytoplasmic core called the rachis through large intercellular bridges (Kimura and Motegi, 2025; Priti et al., 2018); cytoplasmic connectivity between blastomeres in the *Xenopus* embryo (Landesman et al., 2000); the plasmodesmata in plants which are microscopic cytoplasmic channels that traverse the cell walls of plant cells (Bayer and Benitez-Alfonso, 2024); and the germline cysts found across a wide range of species from *Drosophila* to mammals where each germ cell is connected via interconnected bridges (Chaigne and Brunet, 2022). In these contexts, precise regulation of individual cell volume within the shared cytoplasm is important for development, differentiation, and physiological functions.

Our present study focuses on mouse fetal germline cysts, where sister cells derived from a common progenitor remain interconnected through intercellular bridges formed by incomplete cytokinesis (Fig. 1A-B) (Chaigne and Brunet, 2022). A key aspect of germline cyst development, particularly in females, is the highly regulated process of oocyte differentiation, which often involves a dramatic increase in the volume of a few cells. During this process in mice, both cyst structure and fragmentation of germline cysts play critical roles (Ikami et al., 2023). The mouse female germline cysts are not simple linear chains, but rather complex branched structures (Fig. 1A), which are then partially pruned during fragmentation. Future oocytes tend to be connected by more intercellular bridges and become physically larger due to cytoplasmic transfer from neighboring cells, even before cyst fragmentation. Therefore, there is a competitive regulation of cell volume. Active and directional cytoplasmic transport via microtubules with motor proteins was found in *Drosophila* female cysts (Clark et al., 2007; Gutzeit, 1986; Lu et al., 2022). Although such transport was investigated in mice (Lei and Spradling, 2016), it has not been fully characterized.

**Figure 1:**
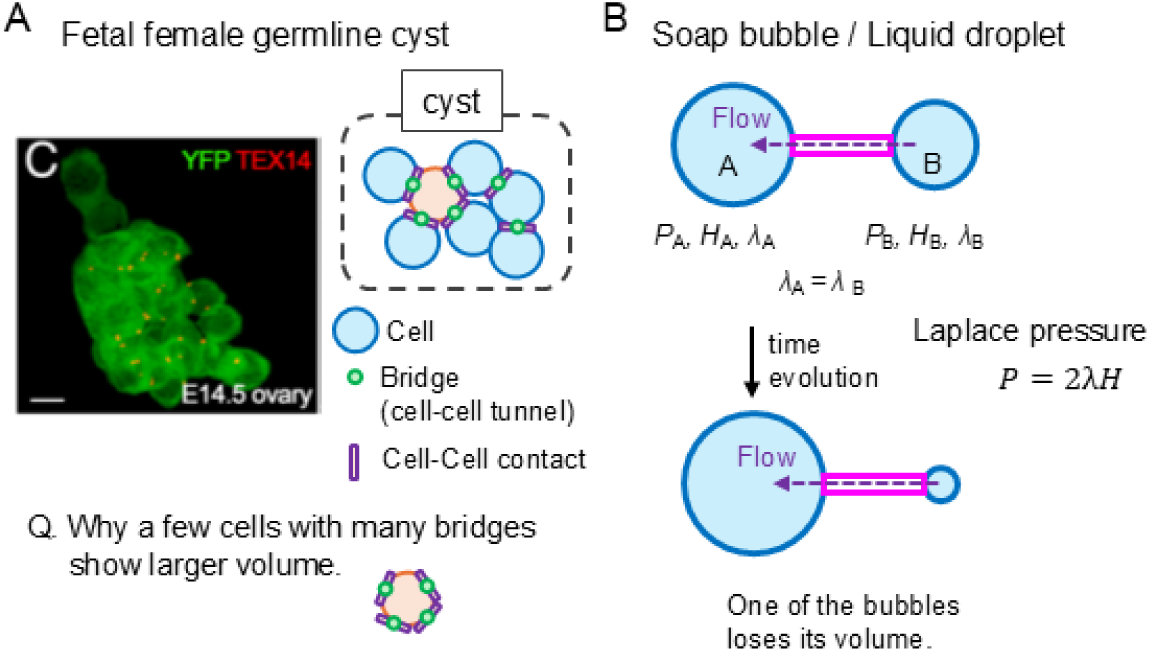
Structures of fetal female germline cysts and mechanical properties of cells. A. Structure of a fetal female cyst. Cells adhere via E-cadherin to generate cell-cell contacts (purple rectangle), where intercellular bridges (green circle) enable cytoplasmic sharing. The microscopic image is reproduced from (Ikami et al., 2023); a single clone is labeled by YFP. YFP, cytoplasm; TEX14, intercellular bridges. B. Soap bubble model. A cell is modeled as a soap bubble or liquid droplet with hydrostatic pressure (*P*), surface tension (*λ*), and curvature (*H*) related by the Laplace pressure equation (*P*=2*λH*). Subscripts denote cell IDs (e.g., *P*_A_ for A-cell). Cytoplasmic flow occurs due to pressure differences between connected cells (e.g., *P*_A_ vs. *P*_B_).

Recently, mechanical forces have been reported to play critical roles in various morphogenetic events and differentiation (Heisenberg and Bellaïche, 2013; Koyama et al., 2023; Kurotaki et al., 2007; Sagner et al., 2012; Savin et al., 2011). Cell volume and shape are also regulated by mechanical forces (Cuvelier et al., 2023; Devany et al., 2023; Farhadifar et al., 2007; Koyama et al., 2012; Miyoshi et al., 2006; Okuda et al., 2015; Runser et al., 2024; Xie et al., 2024). Cytoplasm is flowed by its hydrostatic pressure which is, at least, partially provided from cell surface tensions (Fujita and Onami, 2012; Sedzinski et al., 2011). While active and directional cytoplasmic transport may contribute to oocyte growth, it is plausible that such transport would increase hydrostatic pressure in the larger cells, potentially creating a counteracting force that could offset or reverse cytoplasmic flow. Therefore, a precise control of cytoplasmic pressure may be involved in robust cyst development and the stable establishment of cell fate. In the case of *C. elegans* germ cells connected to the rachis through intercellular bridges as mentioned above, cell surface tensions generate differences in inner pressures, leading to biased cell volume (Chartier et al., 2021). In contrast to the *C. elegans* germ cells, mouse female germ cells exhibit a distinct structure: they lack a rachis, and crucially, the number of intercellular bridges varies among individual cells within the cyst. This complexity necessitates studying how mechanical factors regulate volume under conditions where connectivity is heterogeneous.

Here, we theoretically investigated the contribution of purely mechanical properties of cells to cytoplasmic pressure regulation and subsequent volume distribution within mouse female germline cysts. We developed a simplified model where each cell is analogous to a soap bubble, allowing us to explore the relationship between cell surface tension, surface curvature, and internal cytoplasmic pressure, as described by the Laplace pressure law. Our model also assumes factors, including the number of bridges, cell-cell contact area, cell cycle, and cyst fragmentation as described later. By analyzing this model, we elucidated possible contributions of these cell-intrinsic properties to the biased cell volume distribution in the female germline cyst.

## 1. Theory

### 1-1. Definition of mechanical properties of cells

We developed a theoretical model to investigate the mechanical properties governing cell volume regulation within germline cysts. Figure 1A illustrates the structure of the female fetal germline cysts in mice. During this stage, cells exhibit robust adhesion via E-cadherin proteins at cell-cell contacts, where intercellular bridges enable cytoplasmic sharing (Fig. 1A) (Ikami et al., 2023).

To model the mechanical behavior of these cells, we drew an analogy to a soap bubble, considering cell surface tensions and internal hydrostatic pressures. This simplified model, often employed in cell mechanics due to its experimental basis and inherent simplicity, relates internal hydrostatic pressure (*P*), surface tension (*λ*), and surface curvature (*H*) via the Laplace pressure equation (*P* = 2*λH*) (Fig. 1B) (Sedzinski et al., 2011; Wang et al., 2021). Subscripts for each parameter denote specific cell IDs (e.g., *P*_A_ for A-cell, *P*_B_ for B-cells). A critical assumption of typical soap bubble-based models is that surface tension (*λ*) remains constant regardless of cell size, because the surface tension may be determined by the contractility of the cortical actomyosins rather than by elasticity (Sedzinski et al., 2011). Consequently, as a cell’s volume increases, its surface curvature (*H*) simultaneously decreases. Under the constant *λ* assumption, this leads to a decrease in the pressure (*P*). It is known that in such soap bubble-based systems (and in real soap bubbles), even a slight difference in volume between connected cells is temporally amplified (Sedzinski et al., 2011), leading to the complete loss of volume in one of the cells due to cytoplasmic flow driven by pressure differences (e.g., *P*_A_ vs. *P*_B_). This knowledge implies an instability if not properly regulated.

### 1-2. Assumption of constant cell-cell contact number throughout the cell cycle

On the basis of the above model, we aimed to evaluate the effect of the number of cell-cell contacts with intercellular bridges on cell volume. In the real tissues, the number of cell-cell contacts for each cell should be temporally changeable due to cell division and cyst fragmentation. However, to purely extract the effect of the number of cell-cell contacts, we constructed a model where the number of cell-cell contacts was set to be constant throughout the simulations even when cell division occurred as follows. In Fig. 2A, B-cell has two cell-cell contacts with A_1_- and A_2_-cells, while A_1_- and A_2_-cells have one cell-cell contact. We assumed that the A_2_-cell was equivalent to the A_1_-cell (i.e., the same parameter values are shared). In vivo fetal germ cells undergo multiple rounds of cell division, with each division theoretically reducing the cell volume by 50%. However, the overall cell volume does not decrease during multiple rounds of the cell cycle (Ikami et al., 2023), indicating that concurrent cell growth (i.e., nutrient uptake and synthesis (Devany et al., 2023; Phillips et al., 2008; Wu et al., 2022)) occurs during interphase. Therefore, during a cell cycle, cell volumes increase due to cell growth, while the cytoplasm flows through the bridges due to pressure differences (Fig. 2A, 1^st^ panel). Then, these cells synchronously experience cell division in a manner similar to *in vivo* germline cysts. We assumed a fixed cell division plane so that the connection of the original three cells (B-, A_1_-, and A_2_-cells) is maintained (Fig. 2A, 2^nd^ and 3^rd^ panels). Furthermore, although the proliferated cells are also connected to their sister cells in *in vivo* cysts, we forcibly deleted these cells in our model so that the number of cell-cell contacts in B-, A_1_-, and A_2_-cells remains constant (Fig. 2A, 3^rd^ and 4^th^ panels). Under these settings, combined with the equivalence between A_1_- and A_2_-cells, the system is virtually composed of just two cells with a constant number of cell-cell contacts (Fig. 2A, magenta dashed line boxes). Note that we will also show a full model incorporating all proliferated cells later.

**Figure 2:**
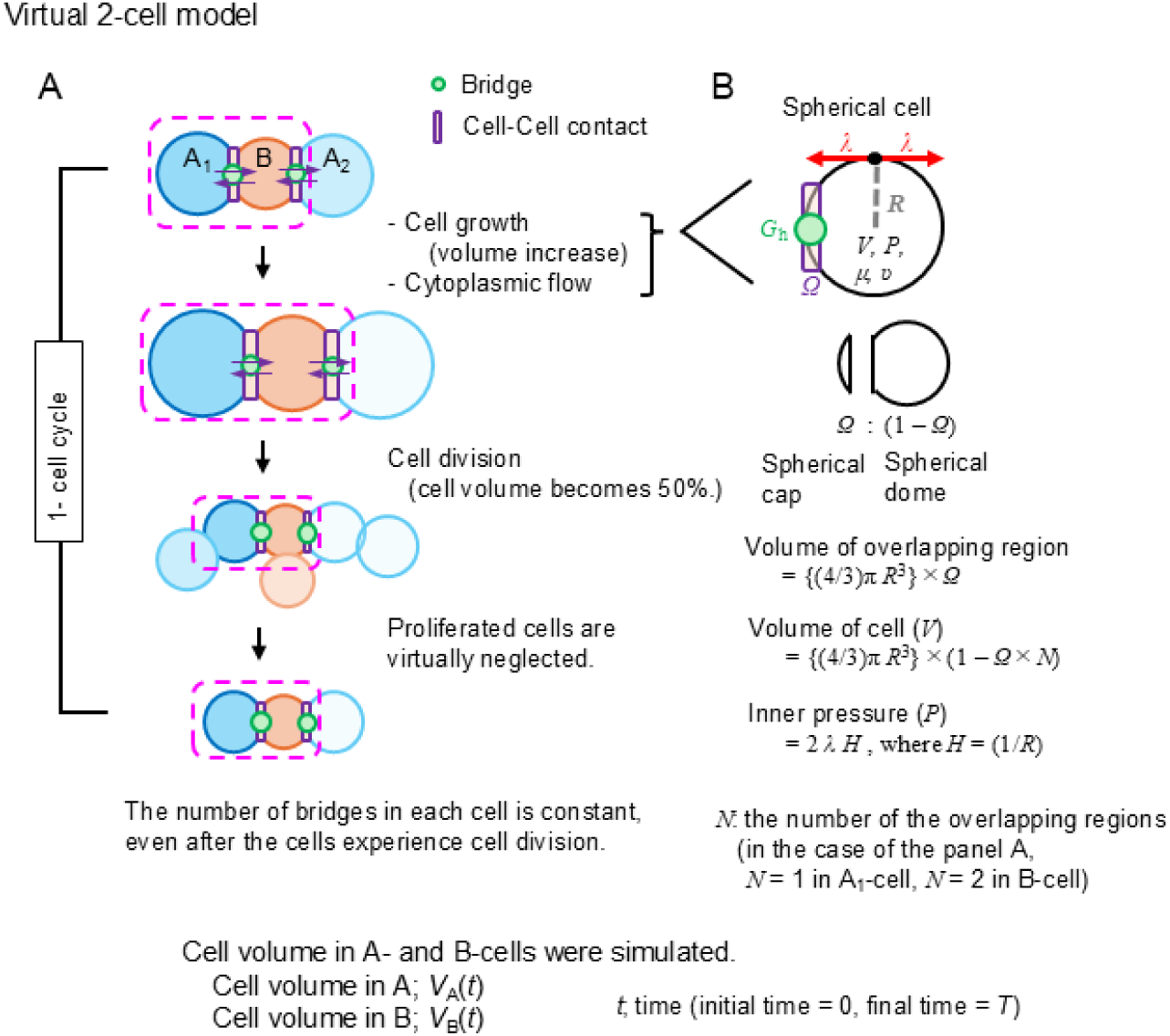
Virtual 2-cell model. A. Modeling of a virtual 2-cell system. The number of cell-cell contacts with intercellular bridges (magenta box) remains constant. One cell cycle includes cell growth, intercellular cytoplasmic flow, cell division, and neglect of proliferated cells. B. Calculation of inner pressure. During cell growth and cytoplasmic flow, inner pressures were calculated based on the Laplace pressure, considering spherical dome shapes due to cell-cell contact regions. Illustrates spherical dome and spherical cap shapes.

### 1-3. Assumption of cell-cell contact regions

Next, we modeled the cell-cell contact region (Fig. 1A). Due to the cell-cell contacts via E-cadherin, the cells show spherical dome shapes (Fig. 2B). In the absence of E-cadherin junctions, the interconnected bridges are formed, but the cell-cell contacts are not. The radius of the spherical dome is denoted as *R*. The ratio of the volume in the spherical dome to that in the sphere whose radius is the same as *R* denotes (1 - *Ω*) (Fig. 2B). When the number of cell-cell contacts (*N*) is larger than 1, the ratio of the volume becomes (1 - *ΩN*). According to these relationships, if the cell volume (*V*) is given, *R* can be calculated by considering *Ω* and *N* (Fig. 2 and S1). Although *Ω* would be complicatedly changed in the *in vivo* tissues, we simply assumed that *Ω* is constant throughout a simulation to extract fundamental behaviors in the presence of the cell-cell contacts.

### 1-4. Calculation of pressure and cytoplasmic flow rate

After the calculation of *R*, we can calculate *P* using the Laplace pressure equation: *P* = 2λ*H* = 2*λ*/*R* (Fig. 2 and S1). The difference in *P* between interconnected cells generates intercellular cytoplasmic flow. The volumetric flow rate through the bridges may be described by the Hagen–Poiseuille equation (Phillips et al., 2008). This equation states that the volumetric flow rate in a cylindrical tube is proportional to both the pressure difference and the fourth power of the tube radius, and inversely proportional to both the tube length and viscosity (*μ*). In our model, we subsumed these geometric factors (i.e., radius, length, and other structural properties) into a single parameter *G*_h_ (“geometry of hole”), leading to the equation of the flow rate as shown in Fig. S1. Both *G*_h_ and *μ* are constant throughout a simulation.

### 1-5. Overall procedure of simulation

Our simulation procedure is as follows (Fig. S1). As the initial condition, the surface tensions are given as constant (*λ*_A_ = *λ*_B_ = const.), and the initial cell volumes (*V*_A_ and *V*_B_) in A_1_- and B-cells are given. After simulations start, the radii of the cells (*R*_A_ and *R*_B_) are calculated according to *Ω* and *N*; the pressures (*P*_A_ and *P*_B_) are calculated by considering the Laplace pressure equation; the intercellular hydraulic cytoplasmic flow rates are calculated; the rates of cell volume changes are determined by the summation of the intercellular cytoplasmic flow rates and cell growth rate which will be defined later; the volume changes are numerically calculated by using the Euler method. Note that the above step involving the Laplace pressure equation was assumed to be a quasi-static process, ensuring that the Laplace law is satisfied at each time step throughout the temporal evolution. After the time period of one cell cycle, the cells are divided, and the proliferated cells are deleted to maintain the virtual 2-cell situation.

In our simulations, the initial *R*_A_ was set to be 1.0 as the characteristic length scale, and the time period of one cell cycle was set to be 1.0. Other default parameter values are *λ* = 1.0, *μ* = 0.2, and *G*_h_ = 0.2, unless noted otherwise.

## 2. Results

### 2-1. Virtual 2-cell system without cell growth and division

#### 2-1-1. The virtual 2-cell system is unstable

We first confirmed the instability of the system composed of interconnected cells, as illustrated in Fig. 1B. We then performed simulations in the absence of cell division. Figure 3A presents simulation results when the overlapping region (*Ω*) between the cell-cell contacts was set to zero (i.e., *Ω* = 0%). The A-cell always had one bridge (*N*_A_=1), while the B-cell had either one (*N*_B_=1) or two bridges (*N*_B_ =2) (Fig. 3A, upper or lower panel, respectively); if *N*_A_=1 and *N*_B_=1, these two cells are equivalent. When the initial volume ratio of B-cell to A-cell (*V*_B_(*t*=0)/*V*_A_(*t*=0)) was exactly 1.0, the cell volumes remained constant over time, even if *N*_B_ > *N*_A_. However, when the B-cell had a slightly larger volume at the initial condition, such as *V*_B_(0)/*V*_A_(0) = 1.01, the B-cell volume was continuously increased, whereas the A-cell volume was decreased, eventually resulting in the complete loss of A-cell volume. This demonstrates that *V*_B_(0)/*V*_A_(0) = 1.0 represents an unstable equilibrium state under these conditions, as we had expected. In other words, *in vivo* germline cysts should have mechanisms for stabilization, which prevent the complete loss of germ cells in the cyst due to a small degree of difference in cell volume. Note that we did not include nuclei in our considerations, which cannot pass through the intercellular bridges.

**Figure 3:**
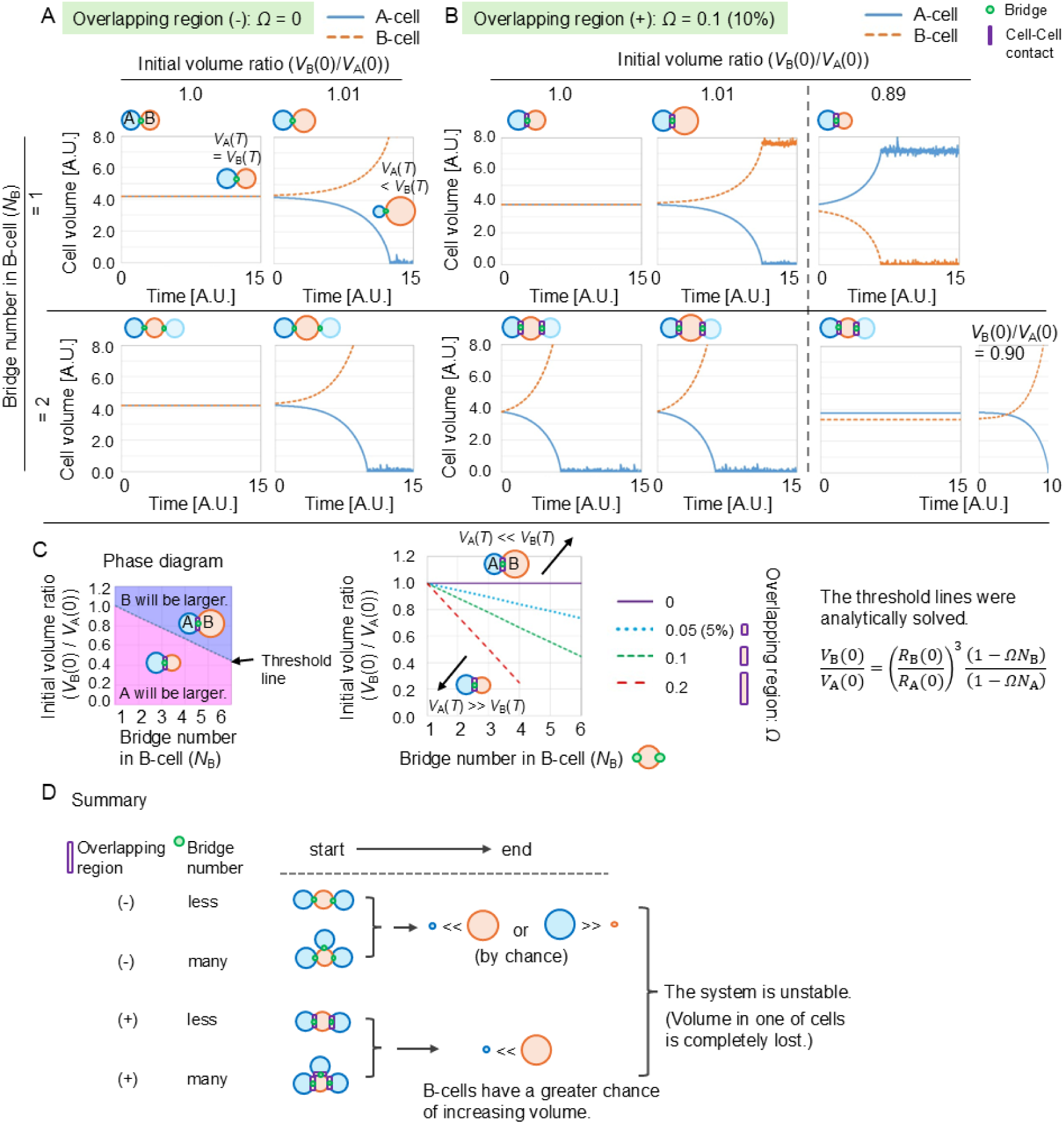
Simulation in virtual 2-cell model without cell division and cell growth. A. Simulation results without overlapping region (*Ω* = 0%). Bridge number in A-cell (*N*_A_) = 1; B-cell (*N*_B_) = 1 or 2. Volume changes for initial ratio *V*_B_(0)/*V*_A_(0) = 1.0 (unstable equilibrium) and 1.01 are shown. B. Simulation results with overlapping region (*Ω* ≠ 0%). Bridge number in A-cell (*N*_A_) = 1; B-cell (*N*_B_) = 1 or 2. Volume changes for initial ratios of 1.0 and ∼0.89 are shown. C. Conditions defining unstable equilibrium states for various *Ω* values. Lines represent thresholds of *V*_B_(0)/*V*_A_(0) for each *Ω* value: above the line, B-cell volume increases (*V*_A_(*T*)<*V*_B_(*T*)); below the line, A-cell volume increases (*V*_A_ (*T*)>*V*_B_(*T*)). D. Summary of the results in the absence of cell cycle. In the presence of the overlapping region, the B-cell is likely to increase its volume, especially when the B-cell has more bridges. In the absence of the overlapping region, the probability of whether the A-cell or B-cell increases its volume is even, and one of the cells completely loses its volume.

#### 2-1-2. Overlapping region changes the condition of unstable equilibrium state

Next, we examined the influence of the overlapping region (*Ω*≠0%) on the instability (Fig. 3B), recapitulating E-cadherin junctions between cyst germ cells. When *N*_B_ =1 (Fig. 3B, upper panels), the simulation results were largely similar to the *Ω* = 0% case (Fig. 3A); an initial ratio of 1.0 led to constant volumes, while a slight deviation caused volume divergence. However, when *N*_B_=2 (Fig. 3B, lower panels), the initial condition *V*_B_(0)/*V*_A_(0) = 1.0 no longer represented an equilibrium state; instead, it resulted in an increase in B-cell volume and a decrease in A-cell volume. Interestingly, a new unstable equilibrium state emerged at approximately *V*_B_(0)/*V*_A_(0) = ∼0.89 (Fig. 3B). If the initial ratio was slightly larger than 0.89 (e.g., *V*_B_(0)/*V*_A_(0) = 0.90), B-cell volume was continuously increased. Conversely, if *N*_B_ =1 and the initial ratio was ∼0.89, the A-cell volume was continuously increased. If *N*_B_ =2 and the initial ratio was less than 0.89, A-cell volume was continuously increased as later shown in Fig. 3C.

We plotted these unstable equilibrium states for various values of *N*_B_ and *V*_B_(0)/*V*_A_(0) (Fig. 3C). The left panel in Fig. 3C shows how to read a phase diagram for a certain value of *Ω*: the unstable equilibrium state is shown as a line that represents the critical threshold determining which cell, A or B, would continuously increase in volume. Conditions above the dotted line led to an increase in B-cell volume (*V*_A_(*t*=*T*) < *V*_B_(*t*=*T*)), while conditions below the dotted line resulted in an increase in A-cell volume (*V*_A_(*T*) > *V*_B_(*T*)), where *T* is the final simulation time. In the middle panel in Fig. 3C, the threshold lines for different values of *Ω* are shown; the threshold lines can be analytically solved by considering the condition satisfying *P*_A_ = *P*_B_ (Fig. 3C, right panel). For larger *Ω* values with *N*_B_ ≥2, the unstable equilibrium states shifted to smaller values of *V*_B_(0)/*V*_A_(0). These results indicate that the B-cell can more easily increase its volume in the presence of numerous bridges and larger overlapping regions (Fig. 3D), highlighting the significant role of cell-cell contact regions in modulating volume dynamics. This can be a candidate mechanism for explaining the biased increase in cell volume in fetal female germline cysts (Fig. 1A), although this model has not yet considered the cell cycle. In addition, the values of the surface tension (*λ*), the geometric effect of the bridges (*G*_h_), and the coefficient of viscosity (*μ*) did not affect the fate of cell volume but did affect the rate of the volume changes (Fig. S2).

### 2-2. Virtual 2-cell system with cell growth and division

#### 2-2-1. Cell growth and division stabilize the system

We next incorporated the cell cycle, including both cell division and cell growth, into our virtual 2-cell model. Typical somatic cells undergo cell growth (driven by nutrient uptake and synthesis) during interphase (Devany et al., 2023; Wu et al., 2022), and subsequently, their volume decreases by 50% due to cell division, as mentioned in the Introduction. We assumed two scenarios for cell growth rates (Fig. S3; constant rate or cell cycle-dependent rate). We will show one of the two scenarios below, because the two scenarios resulted in qualitatively similar simulation outcomes (Fig. S4). Figure 4A shows simulation outcomes under conditions similar to Figure 3B, but with the inclusion of cell growth and division. Note that, as previously mentioned, the cell cycle is synchronized in a cyst. Although the number of cell cycles is limited to about five *in vivo*, we performed simulations over numerous rounds to assess the stability and robustness of the volume distribution. In the absence of the cell-cell contact regions (Fig. 4A; *Ω* = 0%), each cell showed an oscillatory behavior of its volume due to cell growth and division. Importantly, although the initial volume ratio was set to be biased (*V*_B_(0)/*V*_A_(0) = 0.89 ≠ 1.0), the volume in B-cell became equal to that in A-cell, which was contrast to Figure 3 where a slight difference in the initial volume was continuously enhanced (e.g., Fig. 3B, *Ω* = 0%, *V*_B_(0)/*V*_A_(0) = 1.01). Therefore, cell growth and division stabilized the systems, but did not lead to biased cell volume in the absence of cell-cell contact regions.

**Figure 4:**
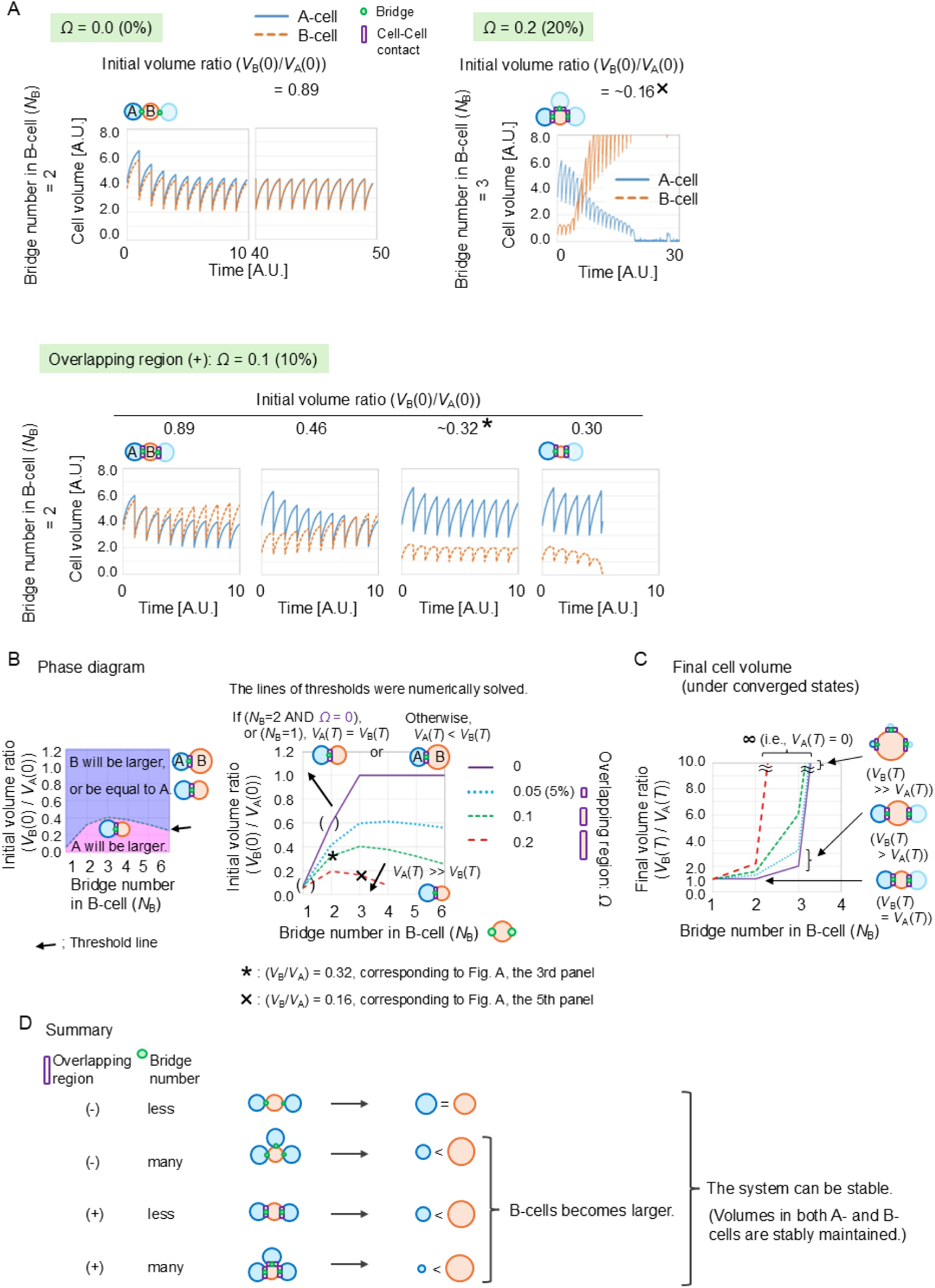
Simulation in virtual 2-cell model with both cell division and cell growth. A. Simulation results with *Ω* = 0, 10, or 20% and *N*_B_=2 or 3 (*N*_A_=1). Volume changes for initial ratio *V*_B_(0)/*V*_A_(0) = ∼0.89 and 0.30 are shown. In the first panel (*Ω* = 0), the simulation result is shown for a long simulation time (*t* = 0∼50). In the bottom panel (*Ω* = 0.1), the behaviors under the different initial conditions (*V*_B_(0)/*V*_A_(0)) are shown. B. Conditions defining threshold values of *V*_B_(0)/*V*_A_(0) for various *Ω* values. For *Ω* > 0, *N*_B_ ≥2: above line, B-cell volume increases; below line, A-cell volume increases. For *Ω* = 0 or *N*_B_=1: above line, *V*_A_(*T*) = *V*_B_(*T*); below line, A-cell volume increases. Asterisk and cross-marked points correspond to conditions in A. C. Final volume ratios (*V*_B_(*T*)/*V*_A_(*T*)) at convergence for various *Ω* values under conditions where B-cell volume increased. Shows specific ratios for *N*_B_=1,2,3 when *Ω* = 0.2. D. Summary of the results in the presence of cell cycle. In the presence of the overlapping region, the B-cell is likely to increase its volume, especially when the B-cell has more bridges. In the absence of the overlapping region, both the A- and B-cells can maintain their volume, and the volume of these cells is equalized if the bridge number in the B-cell is less.

#### 2-2-2. Cell growth and division enhance the likelihood of volume increase in cells with numerous bridges

Next, we evaluated the effect of the overlapping regions of cell-cell contacts in the presence of cell growth and division. In Figure 4A, with *Ω* = 10% and *N*_B_=2 (*N*_A_=1), an initial volume ratio of *V*_B_ (0)/*V*_A_(0) = ∼0.89 led to the B-cell volume becoming larger than the A-cell volume. Therefore, the overlapping regions caused the biased volume increase in B-cell (*Ω* = 10 vs. 0%). With *Ω* = 10%, even if *V*_B_(0)/*V*_A_(0) was quite small (0.46), the B-cell became larger than the A-cell. A new threshold value of *V*_B_(0)/*V*_A_(0) for volume increase/decrease emerged at approximately 0.32, which was much smaller than the threshold value in Fig. 3B (0.89). If the initial ratio was smaller than this threshold (e.g., 0.30), B-cell volume was completely lost. Under the condition of larger *Ω* (20%) with larger *N*_B_ (3), the threshold was further decreased (∼0.16). These results theoretically indicate that, in proliferating germ cells, the cells harboring the numerous bridges with larger overlapping regions have a higher likelihood of volume increase.

#### 2-2-3. Systematic analysis of the likelihood of volume increase in cells with numerous bridges

Then, we systematically analyzed these threshold values of *V*_B_(0)/*V*_A_(0) for volume increase/decrease. We generated a phase diagram (Fig. 4B) in a manner similar to that in Fig. 3C. As explained in the left panel in Fig. 4B, the threshold line was calculated; conditions below the line resulted in an increase in A-cell volume (*V*_A_(*T*) > *V*_B_(*T*)), while conditions above the line led to an increase in the B-cell volume (*V*_A_(*t*=*T*) < *V*_B_(*t*=*T*)) or to equal volumes between B- and A-cells. We calculated the threshold line through numerous trials of simulations (i.e., numerically solved but not analytically). The right panel in Fig. 4B is the phase diagram for different values of *Ω*. For [*N*_B_ = 2 AND *Ω* = 0] or *N*_B_ = 1 (even with *Ω* > 0), conditions above the lines resulted in the equal volumes between A- and B-cells (*V*_A_ (*T*)=*V*_B_(*T*)). Otherwise, conditions above the lines led to an increase in B-cell volume (*V*_A_(*T*) < *V*_B_(*T*)). Conditions below the lines resulted in an increase in A-cell volume (*V*_A_(*T*) > *V*_B_(*T*)). The asterisk-marked point on the green dashed line in Figure 4B corresponds to the condition in Figure 4A (*V*_B_(0)/ *V*_A_(0) = ∼0.32 with *Ω* = 0.1), and the cross-marked point adjacent to the red dashed line corresponds to *V*_B_(0)/ *V*_A_(0) = ∼0.16 with *Ω* = 0.2. In comparison with Fig. 3C, the threshold lines were largely shifted downward in the phase diagram, indicating that the likelihood of volume increase in the B-cell with numerous bridges was enhanced in the presence of cell growth and division. In other words, even if the initial volume in the B-cell is much smaller by chance, the B-cell can selectively increase its volume through the cell cycle. This mechanism may contribute to robustly selecting cells whose volumes should be increased.

#### 2-2-4. Calculation of cell volume at converged state

Finally, we ran the simulations until they reached converged states, by which we obtained the final cell volume ratio (*V*_B_(*T*)/*V*_A_(*T*)). We focused on the conditions where the B-cell volume increased (i.e., above the threshold lines in Figure 4B). For a certain value of *Ω* and *N*_B_, the final volume ratios remained constant regardless of the initial volume ratios (*V*_B_(0)/*V*_A_(0)), as long as they were above the respective threshold lines. For instance, when *Ω* = 0.2 (20%), the final volume ratio was 1.0 for *N*_B_ = 1 or approximately 2.0 for *N*_B_ = 2 (Fig. 4C, red broken line, *N*_B_ = 1 or = 2). Notably, for *N*_B_ = 3, the ratio became infinite (indicated by ∞), because of the complete loss of the A-cell volume, as also suggested by the panel of Figure 4A, *Ω* = 0.2 (20%). As *Ω* was set to be larger (Fig. 4C; *Ω* = 0.2 vs. 0.1 (green broken line), 0.05 (light blue broken line), or 0 (purple line)), the final volume ratios became larger. Similarly, as *N*_B_ was set to be larger, the final volume ratios became larger. These results indicate that both the bridge number (*N*_B_) and its overlapping degree (*Ω*) contribute to biased cell volume in specific cells (Fig. 4D).

#### 2-2-5. Summary of virtual 2-cell system simulations

As illustrated in Fig. 4D, both the number of bridges and the overlapping region are critical for biased cell volume expansion. This interplay provides a potential mechanism for the asymmetric volume distribution observed in female fetal germline cysts. In contrast, incorporating the cell cycle leads to an equalized volume distribution in systems with fewer bridges and no overlapping regions—a configuration that may reflect the characteristics of adult male germline cysts, as further detailed in the Discussion.

### 2-3. Full model considering both cyst formation and cyst fragmentation

#### 2-3-1. Definition of Full Model

To investigate cell volume regulation in a more complex and biologically realistic setting, we extended our previous model (the virtual 2-cell model) to incorporates stochastic processes observed in mouse germline development. *In vivo* germline cysts exhibit several characteristic features: the cell-cell connection pattern such as branching varies among the cysts, the cell number in a cyst varies, and the intercellular bridge number can be temporally increased due to cell division or decreased due to bridge severing. In addition to both the Laplace pressure and cell cycle in the virtual 2-cell model, our new model assumed both the cyst formation and cyst fragmentation (bridge severing), by which we evaluated the effect of these features on cell volume.

As depicted in Figure 5A, which illustrates cyst formation, the cell division plane was stochastically set (Fig. S5A), leading to various cell-cell connection patterns; i.e., the proliferated cells were not deleted in this full model. Subsequently, the intercellular bridges are stochastically severed, yielding multiple cysts with different cell numbers (Fig. 5A, bottom panel). The resulting cysts then proceed through the next cell cycle, undergoing processes such as cell growth and cytoplasmic flow as defined in our virtual 2-cell model. For the bridge severing process, we defined two models (Fig. S5B): model-#1, biased severing according to bridge number, and model-#2, random severing.

**Figure 5:**
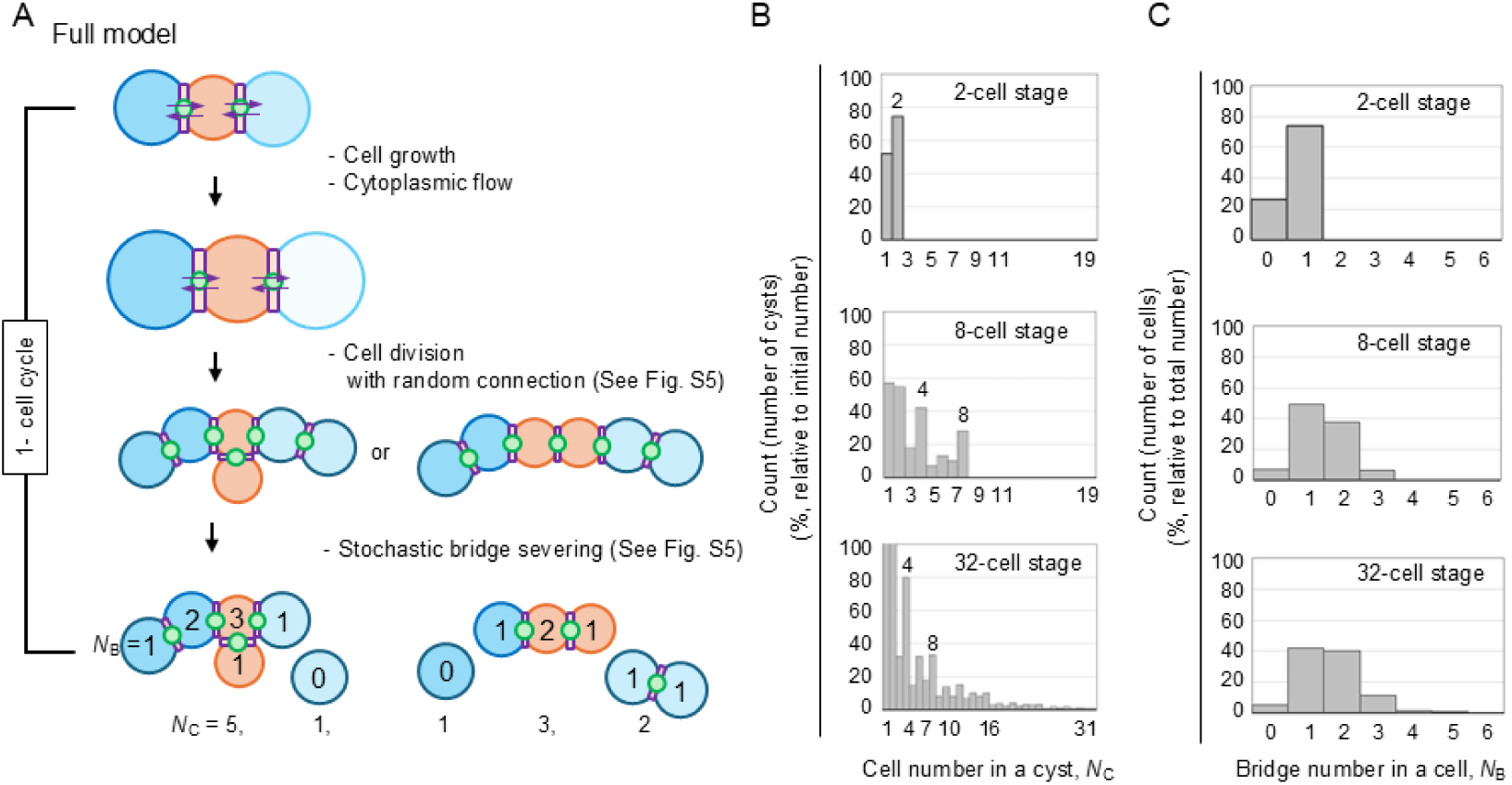
Definition of the full model. A. Schematic of the full model. Connection patterns are randomly determined after cell division, and intercellular bridges are stochastically severed, leading to varied cyst structures before the next cell cycle. B. Histograms of cell numbers within cysts at 2-cell (1 cycle), 8-cell (3 cycles), and 32-cell (5 cycles) stages. C. Histogram of bridge numbers per cell for each stage.

#### 2-3-2. Setting of probability of severing

First, we searched for the probability of bridge severing, which can recapitulate the process of cyst fragmentation *in vivo*. In *in vivo* female germline cysts (Ikami et al., 2023), after starting from cysts composed of 1 cell, the populations of the cysts composed of 4 and 8 cells were enriched after 2 or 3 cell cycles, and then, the number of cells in a cyst diverges from 1 cell to several tens of cells after an addition few cell cycles. Basically, there is a decline in the population of cysts with an increasing number of constituent cells. In our simulations, we set various values of the severing probability for the model-#1 and -#2 (Fig. S6A), and found an optimal value for each model, which can approximate the overall patterns of the populations observed *in vivo* (Fig. S6B). Figure 5B shows the populations under the optimal value in the model-#1 (based on biased severing). Cysts composed of 4 and 8 cells were enriched at the 8-cell stage (3 cell cycles). At the 32-cell stage (5 cell cycles), cysts composed of 4 and 8 cells were still enriched, while cysts with several tens of cells showed smaller populations. The model-#2 (based on random severing) was also able to yield the similar population patterns (Fig. S6A-B). The random severing during cyst formation is sufficient to recapitulate the *in vivo* phenomenon during cyst formation. Following cyst formation, as we previously reported, cell division stops and cyst fragmentation continues, with biased severing occurring throughout this process (Ikami et al., 2023).

We next checked the bridge number in a cell. In *in vivo* female germline cysts (Ikami et al., 2023), the populations of cells with 1 and 2 bridges were enriched, and the populations of cells with 0 or >=3 bridges exhibited lower. Under the condition of the severing probability in Fig. 5B, our simulations enriched the populations of cells with 1 and 2 bridges (Fig. 5C and S6C). Together with Fig. 5B, our model successfully approximated the structures of the cysts during cyst formation from the viewpoint of both the bridge numbers in a cell and the cell number in a cyst. Note that, as we will demonstrate in the following section, the probability of bridge severing itself does not significantly affect cell volume during cyst formation, suggesting that the precise population distribution shown here is not central to our core findings on volume regulation.

#### 2-3-3. Effect of overlapping region and bridge number on cell volume in the full model

We next used the full model to investigate whether both the overlapping regions of cell-cell contacts and the bridge number influence individual cell volumes within the cysts. Figure 6A-i shows the average cell volumes for the cells harboring different numbers of bridges. In the absence of the overlapping region (left upper panel; *Ω* = 0.0), the cell volumes were not significantly different among the cells with different numbers of bridges (*N*_B_). On the other hand, in the presence of the overlapping regions (right upper panel; *Ω* = 10.0%), the cell volumes became larger as the bridge numbers were increased. These trends were also observed under the condition of the bridge severing probability = 0.0 (*P*_0_ = 0.0, lower two panels; i.e., no severing occurs). Similar results were also obtained in the model-#2 (Fig. S7). In Fig. 6A-ii, we quantified the effect of *Ω* on cell volume. Both *Ω* and *N*_B_ positively contributed to the increase in cell volume. Fig. 6A-iii shows the effect of *P*_0_, where *P*_0_ did not significantly affect the cell volume. In conclusion, the positive effects of both the overlapping region and the bridge number were detected in both the full model (Fig. 6) and the virtual 2-cell model (Fig. 4). In addition, the bridge severing probability had no effect on cell volume.

**Figure 6:**
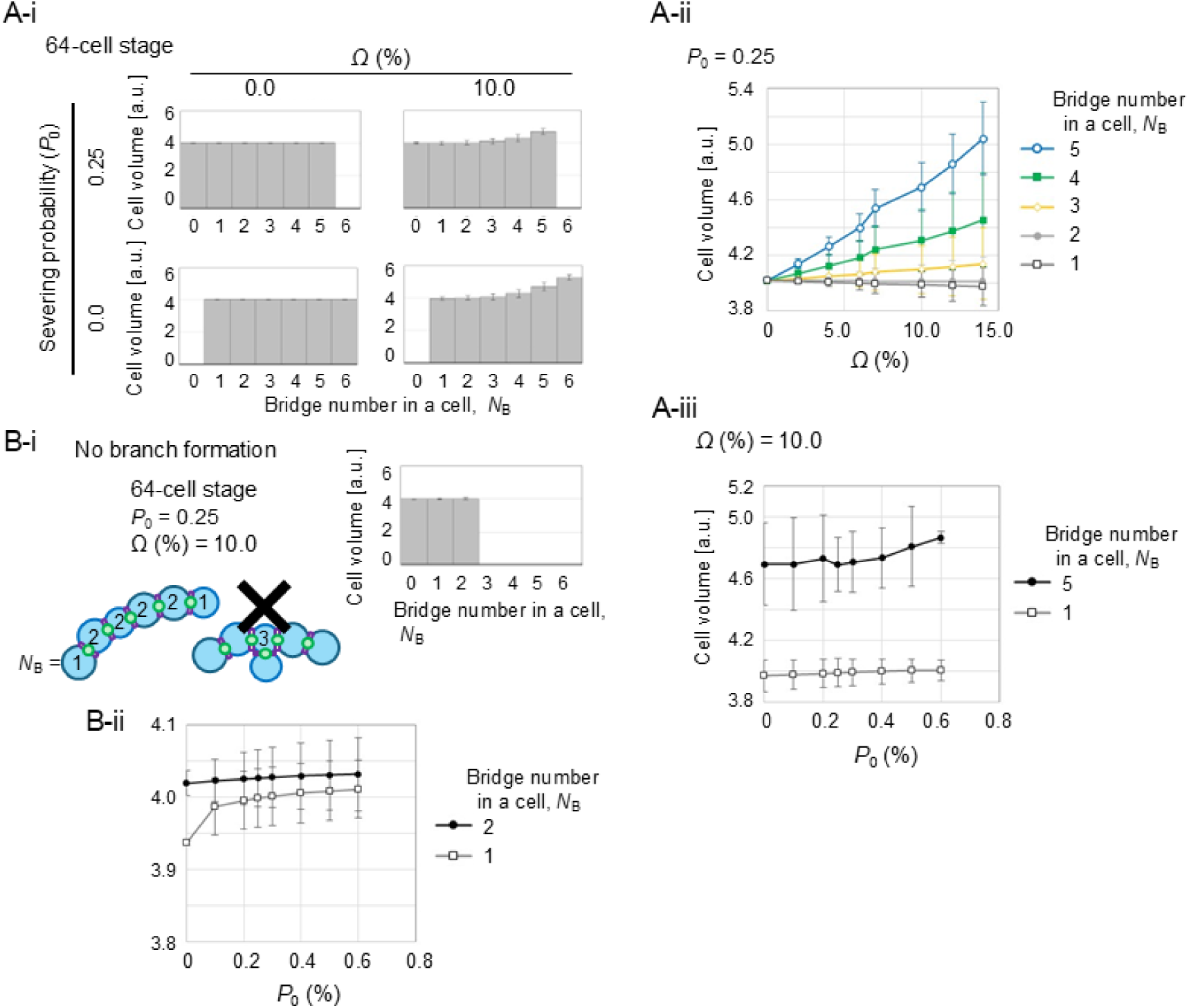
Factors affecting cell volume in the full model. A-i. Cell volumes for cells with different bridge numbers under varying *Ω* and *P*_0_ conditions. A-ii. Effect of *Ω* on the volume of cells with the different number of the bridges, showing specific volume increase for cells with more bridges at larger *Ω*. A-iii. Effect of *P*_0_ on cell volume, indicating no significant changes with varying *P*_0_. B-i. Linear cyst model where no branches are formed, limiting bridge number per cell to less than 3. The histogram was obtained at the 64-cell stage under *Ω* = 10% and *P*_0_ = 0.25. B-ii. Cell volumes in unbranched cysts. Compares volumes of cells with 1 versus 2 bridges, showing limited volume increase even with the maximum bridge count in linear structures.

In the above simulations, the parameter values are identical to those in the virtual 2-cell model, except for the cell growth rate, which was adjusted to almost maintain the total volume in a cyst across the multiple cell cycles (as described in Fig. S3) and *μ* = 0.15.

#### 2-3-4. Branching is effective to cell volume bias

Then, to assess the impact of branching, we modified the full model so that no branches were formed; strictly linear cyst structures were formed solely (Fig. 6B-i). In such linear cysts, the maximum bridge number in a cell is inherently limited to two, as indicated by our simulations (Fig. 6B-i, right panel). Figure 6B-ii shows the cell volumes in these unbranched cysts. In the presence of the overlapping region (*Ω* = 10.0%), even in the cells with the maximum possible bridge number (*N*_B_ = 2), the cell volumes were just slightly increased (approximately ∼4.03) compared to the cells with 1 bridge. This contrasts with the results from the full model with branching (e.g., Figure 6A-ii, where the cells with 5 bridges reached a volume of approximately ∼4.6 when *Ω* = 10.0%). This comparison strongly suggests that the ability to form the branched structures, which allows the cells to acquire a larger number of intercellular bridges, is crucial for establishing pronounced volume biases.

## 3. Impact of cell-cell contacts without bridges on cell volume

In the *in vivo* germline cysts, the cells aggregate regardless of the presence of intercellular bridges, forming tightly packed clusters. (Ikami et al., 2023). At the cell-cell contacts without the bridges, cytoplasm is not directly shared between the two cells. Here, we investigated the effect of such cell-cell contacts on cell volume regulation. Figure 7A illustrates the modified virtual 2-cell model used for this analysis. Similar to the previous simulations in Fig. 4, A-cell and B-cell were initially assumed to have 1 and 2 cell-cell contacts with bridges, respectively (Fig. 7A, “Default”). We then introduced additional cell-cell contacts to the B-cell, either with or without the bridges (Fig. 7A, “Contact number (*N*_new_) = 0, 1, 2…”). For example, the B-cells in the rightmost panels show scenarios where B-cells have a total of 4 cell-cell contacts: either all 4 contacts harbor a bridge (upper panel), or 2 contacts harbor a bridge and the remaining 2 contacts do not (lower panel). This setup allowed us to directly compare the impact of bridge-containing versus bridgeless contacts.

**Figure 7:**
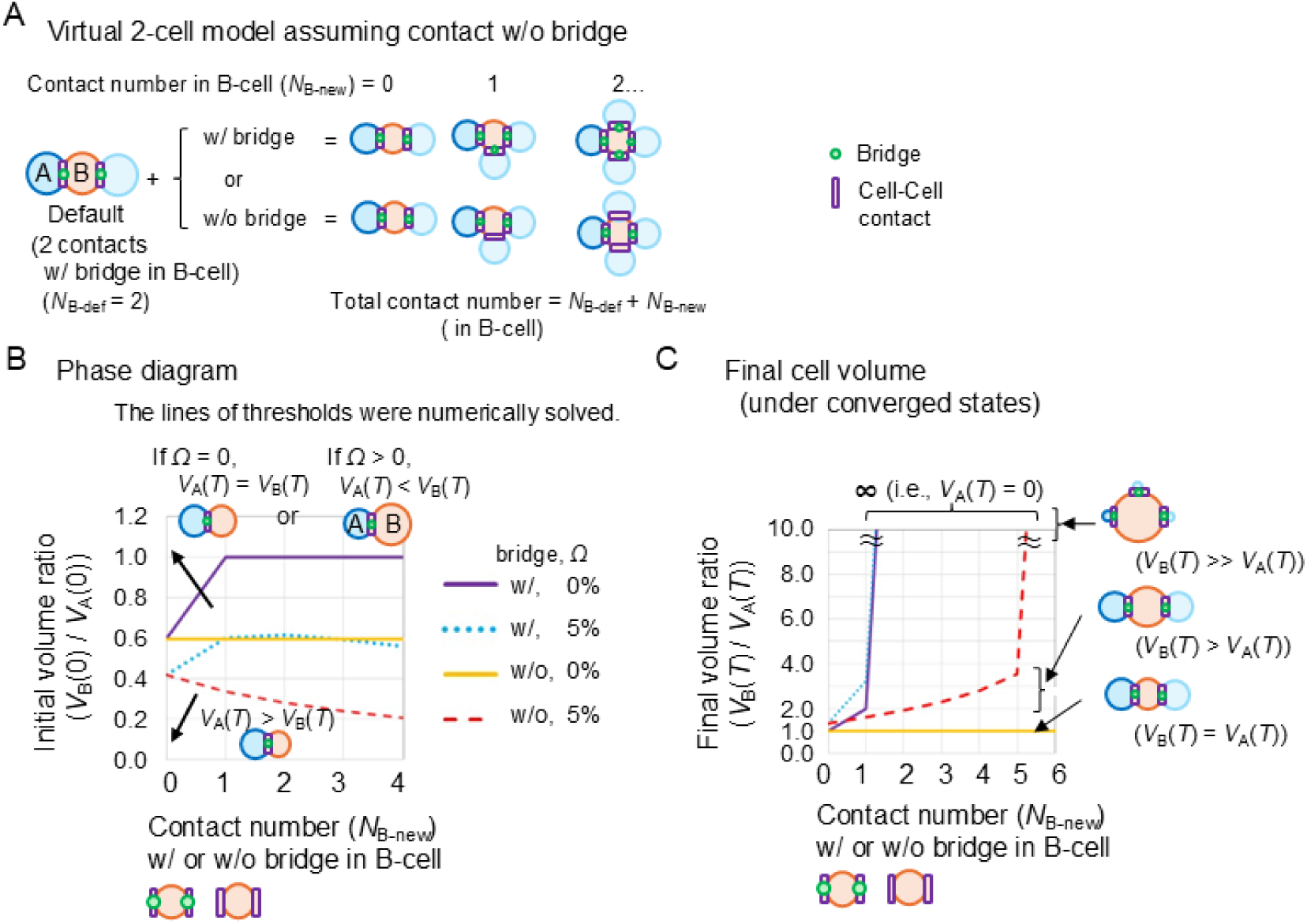
Effect of cell-cell contact without bridges on cell volume. A. Virtual 2-cell model assuming contacts with or without bridges. This panel illustrates scenarios where B-cell has additional contacts, some with bridges and some without. See text for details. B. Threshold values of *V*_B_(0)/*V*_A_(0) for volume change, similar to Figure 4B, considering contacts with/without bridges. For *Ω* > 0 (i.e., 5%; light blue and red lines): above line, B-cell volume increases; below line, A-cell volume increases. For *Ω* = 0 (purple and yellow lines): above line, *V*_A_(*T*) = *V*_B_(*T*); below line, A-cell volume increases. C. Final converged volume ratios (*V*_B_(*T*)/*V*_A_(*T*)) similar to Figure 4C, considering contacts with/without bridges. The legend for the lines is shown in B. Cells harboring contacts without bridges increase the final volume ratio (yellow and red lines), but at a slower rate than those harboring contacts with bridges (purple and light blue lines).

Figure 7B presents the threshold values of the initial volume ratio (*V*_B_(0)/*V*_A_(0)) that determine the direction of volume change, similar to Figure 4B, but now incorporating cell-cell contacts without bridges. With *Ω* > 0, conditions above the lines resulted in B-cell volume increasing (i.e., the final volume ratio will be *V*_A_(*T*) < *V*_B_(*T*)), while conditions below the lines led to A-cell volume increasing (*V*_A_(*T*) > *V*_B_(*T*)). With *Ω* = 0, conditions above the lines resulted in *V*_A_(*T*) = *V*_B_(*T*), whereas below the lines, A-cell volume increased. In comparison between *Ω* = 0 and =5.0%, the presence of the cell-cell contacts, even without the bridges, still influenced the initial conditions that determine the eventual winner of the volume competition (Fig. 7B, yellow vs. red broken line), although the specific thresholds differ from the cases with only the bridge-containing contacts.

Figure 7C shows the final converged volume ratios (*V*_B_(*T*)/*V*_A_(*T*)) under the conditions where the B-cell volume increased, similar to Figure 4C. We observed that the final volume ratios increased as the number of the cell-cell contacts without bridges increased, although the rate of this increase was lower compared to the situations where the additional contacts contained bridges. This finding raises the possibility that the aggregation of the cells in germline cysts contributes to a biased volume increase via cell-cell contacts.

## Discussion

In the present study, we theoretically investigated the mechanism of the competitive regulation of cell volume in mouse female germline cysts. On the basis of cellular mechanics including the cell surface tension and hydrostatic pressure, we constructed mathematical models where the cells shared their cytoplasm through cell-cell contacts with intercellular bridges. In our models, the cells with a larger number of bridges selectively increased their cell volumes, when the cells are tightly contacted with each other (i.e., larger value of overlapping region *Ω*). Furthermore, cell cycle (i.e., cell growth and cell division) contributes to biased volume increase in the cells with more bridges. This bridge number-dependent mechanism accompanying tight cell-cell contacts can be a candidate to explain the biased cell volume observed in the female germline cysts, leading to oocyte differentiation. Our present study theoretically proposes mechanical bases for competitive volume regulation and for robust maintenance of volume in the female germline cysts.

In addition to the above critical factors (i.e., overlapping region, bridge number, and cell cycle (cell growth and cell division)), we identified several factors contributing to the biased cell volume: branching, cell surface tension, cross-section area of the bridge (i.e., diameter of the bridge), and viscosity of cytoplasm. The branching process through cell division leads to an increase in the bridge number, and therefore, is critical for yielding the biased cell volume distributions (Fig. 6). The cell surface tension provides the hydrostatic pressure, which can induce intercellular cytoplasmic flow. The rate of the intercellular cytoplasmic flow is increased according to the geometry of the bridge (*G*_h_) including the radius or inversely proportional to the viscosity of cytoplasm (*μ*) (Fig. S2). By considering these factors, our present study provides experimentally testable hypotheses.

Female gametogenesis proceeds through an initial phase of simultaneous cyst formation and fragmentation, followed by a phase where only fragmentation continues (Fig. 8A). In our present model, we focus on the cyst formation stage (embryonic days 10–14, E10-14), when cell division and cyst fragmentation take place. Our previous work, which focused on the stage following cyst formation stage (E14–P0 (postnatal day 0)), provided evidence for biased bridge severing and variability in cell volume among cells within a cyst. Specifically, cells with a greater number of bridges preferentially exhibited larger volumes. Here, we suggest that biased cell volumes begin to emerge during cyst formation, consistent with our earlier observation that differences in cell volume are first detected at E14.5 (Ikami et al., 2023). The small differences in cell volume observed at E14.5 are likely amplified after cyst formation, leading to larger volume disparities among cells.

**Figure 8:**
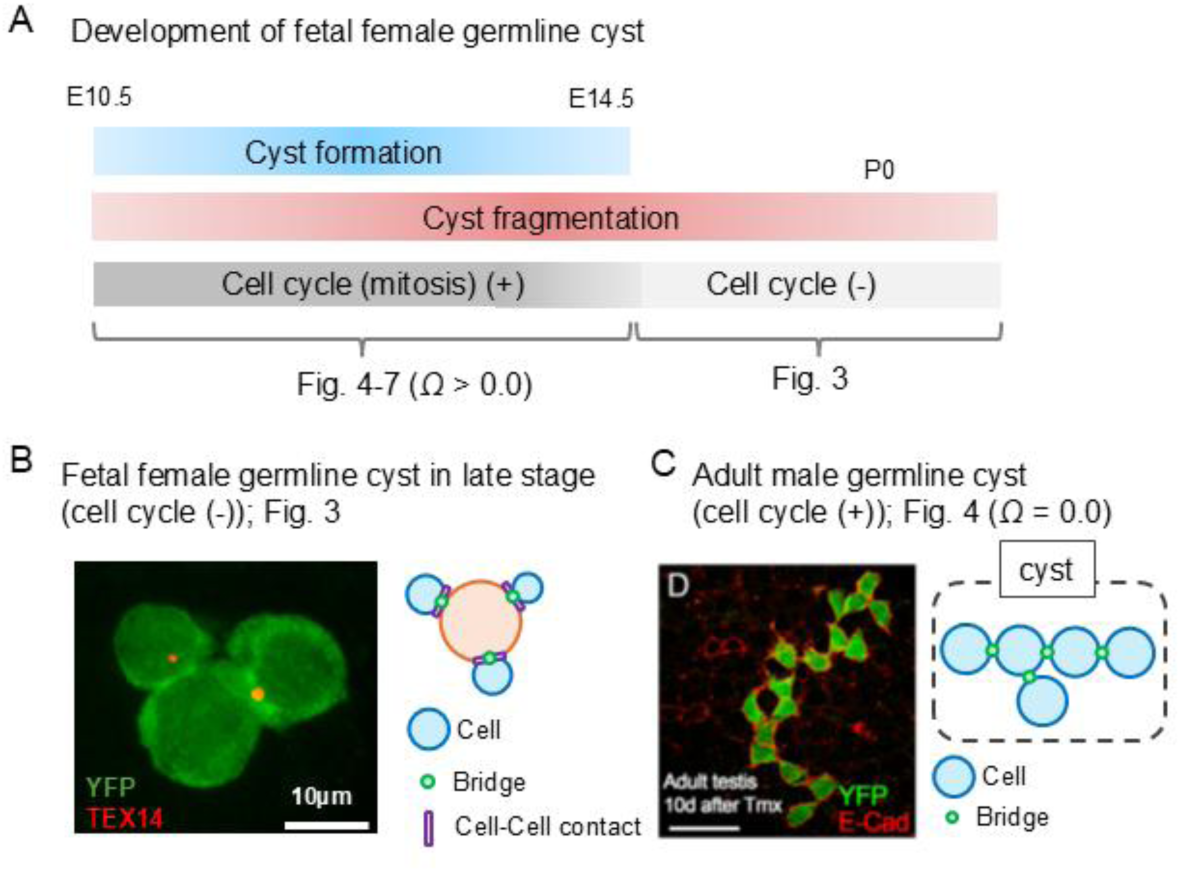
Other type of germline cysts A. Schematic representation of developmental stages in female germ cells. See text for details. B. Structure of a fetal female cyst at later stage. The cells do not undergo cell cycle, and a smaller population of the cells shows extreme increase in volume, leading to oocyte formation. The microscopic image was acquired according to the procedures described previously (Ikami et al., 2023). YFP, cytoplasm; TEX14, intercellular bridges. C. Structure of an adult male cyst before meiosis. Cells exhibit weaker adhesion, but intercellular bridges are present. The cells undergo cell cycle. The microscopic image is reproduced from (Ikami et al., 2023). YFP, cytoplasm; E-cad, E-cadherin.

In the *Drosophila* germline cysts, microtubule-mediated directional transport of cytoplasm is observed, resulting in a biased cell volume (Clark et al., 2007; Gutzeit, 1986; Lu et al., 2022). Our present mechanical model is not incompatible with directional transport-based mechanisms. Rather, our model can support the directional transport-based mechanisms to stably and robustly achieve the biased cell size distribution. In humans, the fetal germline cysts are likely to have a branched structure (Gondos, 1987), implying that the proposed hydrostatic pressure-dependent mechanism may also be applicable to these systems.

Both hydrostatic pressures and surface tensions in cells/tissues are involved not only in phenomena of cell biology such as cytokinesis, blebbing, etc. (Sedzinski et al., 2011), but also in phenomena of tissue morphogenesis such as the blastocyst formation in mice, the behaviors of spheroids (Chan et al., 2019; Dumortier et al., 2019; Mombach et al., 2005). Regulation of these mechanical parameters can stabilize or unstabilize the systems, leading to their robust behaviors and morphological changes, respectively. Our theoretical framework provides mechanical insights into various phenomena observed during germ cell development. We have focused primarily on the dividing female germline cysts (Fig. 1A and 8A). At a subsequent later stage, these cells stop dividing (Fig. 8A) and an extremely biased cell volume increase occurs to form oocytes (Fig. 8B). In consistent with this stage, the results shown in Fig. 3—simulated in the absence of cell cycle—also demonstrated an extreme volume increase in cells with numerous bridges. In contrast, adult male germline cysts undergo mitosis before entering meiosis, and their cell volumes are robustly maintained despite the presence of the intercellular bridges, leading to the proper male germ cell development (Fig. 8C). This scenario is consistent with our simulation in Fig. 4A, which showed that the cell volumes were not biased when cell cycle was assumed but without tight cell-cell contacts. We do not rule out other possibilities, such as a blockage of cytoplasmic flow caused by the small diameter of the bridges, etc. Taken together, our framework demonstrates that the combination of two key mechanical parameters—cell-cell contacts and cell cycle (cell growth and division)—allows the systems to exhibit these three distinct biological states.

Finally, we raise the limitations of our model. Although we assumed that the cells exhibit spherical shapes including spherical domes, the real cell shapes are not necessarily so spherical. The cells within adult male germline cysts adhere to the surrounding tissues, leading to distorted cell shapes. Therefore, the regulation of the hydrostatic pressures becomes more complicated than in our theoretical model. A similar argument applies to other syncytial cells observed in the *C. elegans* gonad. In the context of the *C. elegans* gonad, Chartier et al. incorporated its complicated geometry into their model and demonstrated that mechanical instability, derived from the surface tension and hydrostatic pressure, is critical for regulating biased cell volume (Chartier et al., 2021). Therefore, in order to strictly reproduce the *in vivo* situations, more complicated modeling will be required, while our present model for cytoplasm-sharing cells can be the basic framework.

## Source

The virtual 2-cell model was implemented on the excel software. The full model was written in C language. These are uploaded on figshare as follows (10.6084/m9.figshare.31148572).

## Authorship contributions

Hiroshi Koyama: Writing – review & editing, Writing – original draft, Visualization, Validation, Project administration, Methodology, Investigation, Funding acquisition, Formal analysis, Data curation, Conceptualization. Kanako Ikami: Writing – review & editing, Investigation, Experiment, Conceptualization. Lei Lei: Writing – review & editing, Conceptualization. Toshihiko Fujimori: Writing – review & editing, Conceptualization.

