## supplementary figures for "Mechanically competitive regulation of cell volume in cytoplasm-sharing cells connected by intercellular bridges"

Title

Koyama et al.

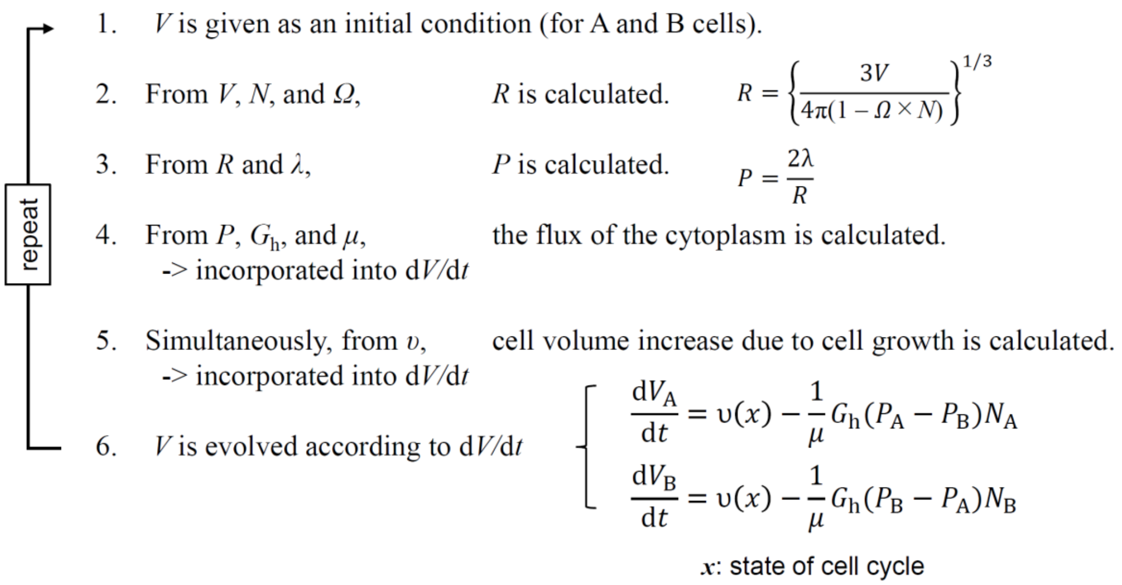


Figure S1 (related to Fig. 2): Simulation procedure of virtual 2-cell model. See text in details. $\upsilon(x)$ is the rate of cell volume increase, which depends on cell cycle as defined in Fig. S3. $x$ denotes the state of cell cycle as defined in Fig. S3.

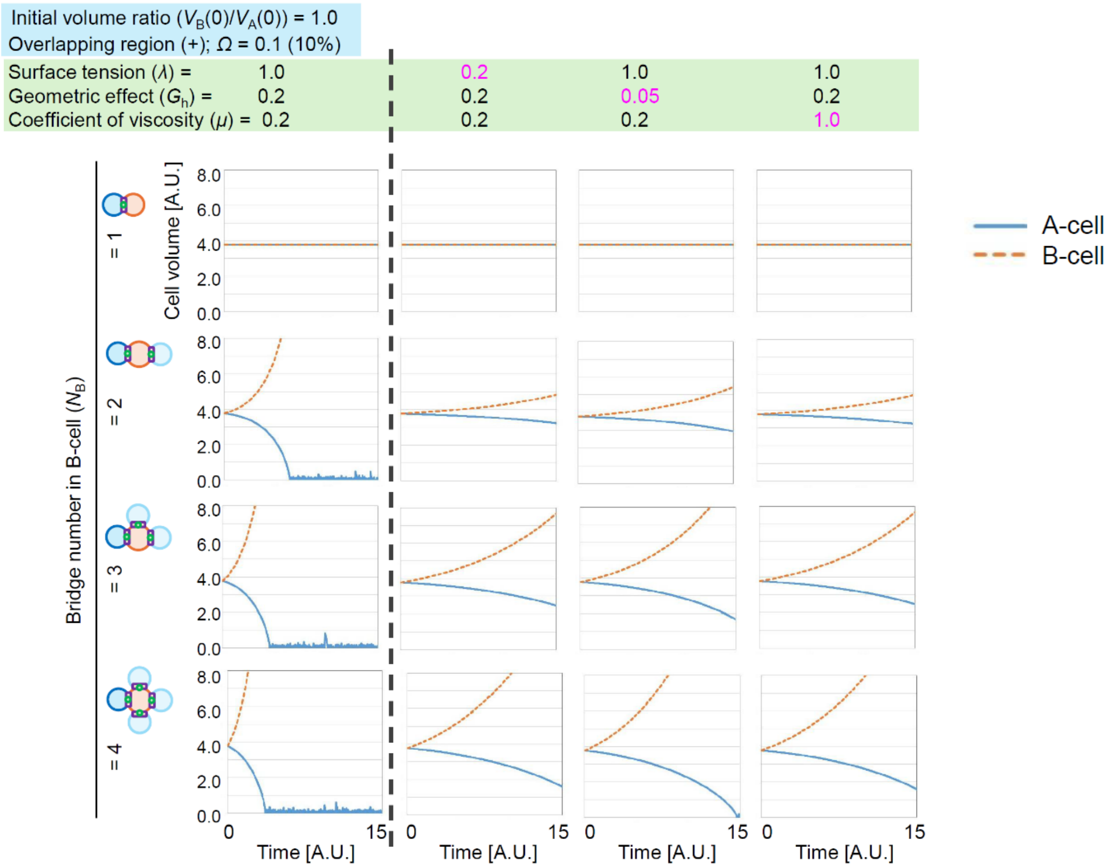


Figure S2 (related to Fig. 3): Simulation outcomes in virtual 2-cell model without cell cycle under various parameter conditions. In this figure, the initial volume ratio and the overlapping region were fixed: $V_B(0)/V_A(0) = 10.0$, $\Omega = 0.1$. The surface tension ($\lambda$), the geometric effect of the bridges ($G_h$) including their cross-sectional area, and the coefficient of viscosity ($\mu$) were varied. $N_B$ was set to be 1~4.

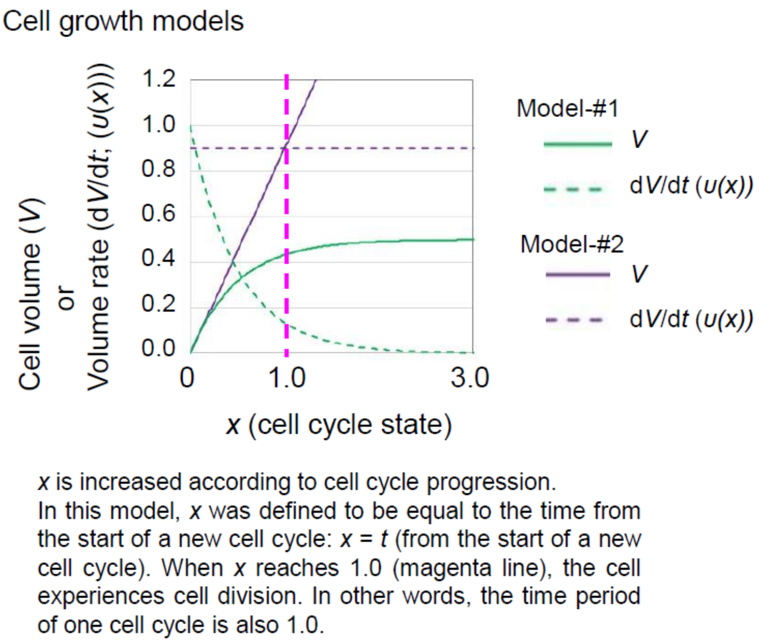


Figure S3 (related to Fig. 4): Definition of cell growth model in virtual 2-cell model with cell cycle. $x$ is the state of cell cycle as defined in the panel. In Model-#1, the rate of cell volume increase was set to be decreased as cell cycle proceeds (d$V$/d$t$; green dot line), leading to a plateau of the volume ($V$; green line). The rate was defined as follows; d$V$/d$t$ = $\varepsilon$ exp(-2$x$) = $\varepsilon$ exp(-2$t$), where $t$ is the time from the start of a new cell cycle, and $\varepsilon$ is the coefficient. $\varepsilon$ is = 1.0 in this figure, whereas $\varepsilon$ = 5.0 in Fig. 4 and 7 and $\varepsilon$ = 9.2 in Fig. 5 and 6. In Model-#2, the rate of cell volume increase was set to be constant (purple dot line), leading to a continuous increase of the volume (purple line) in a manner independent to the cell cycle state.

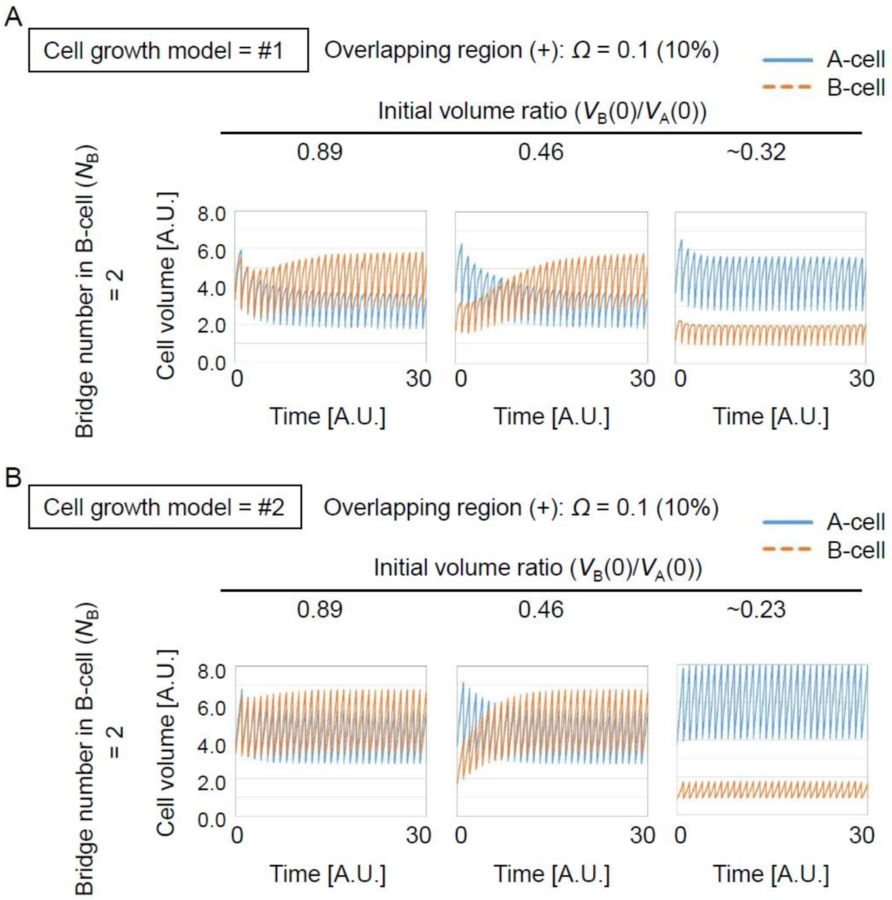


Figure S4 (related to Fig. 4): Simulation outcomes in virtual 2-cell model with cell cycle and cell division in Model-#1 and #2 which were defined in Fig. S3. A. Simulation results in Model-#1 under different values of the initial volume ratio ($V_B(0)/V_A(0)$). B. Simulation results in Model-#1 under different values of the initial volume ratio ($V_B(0)/V_A(0)$). Other parameter values are shown in the panels.

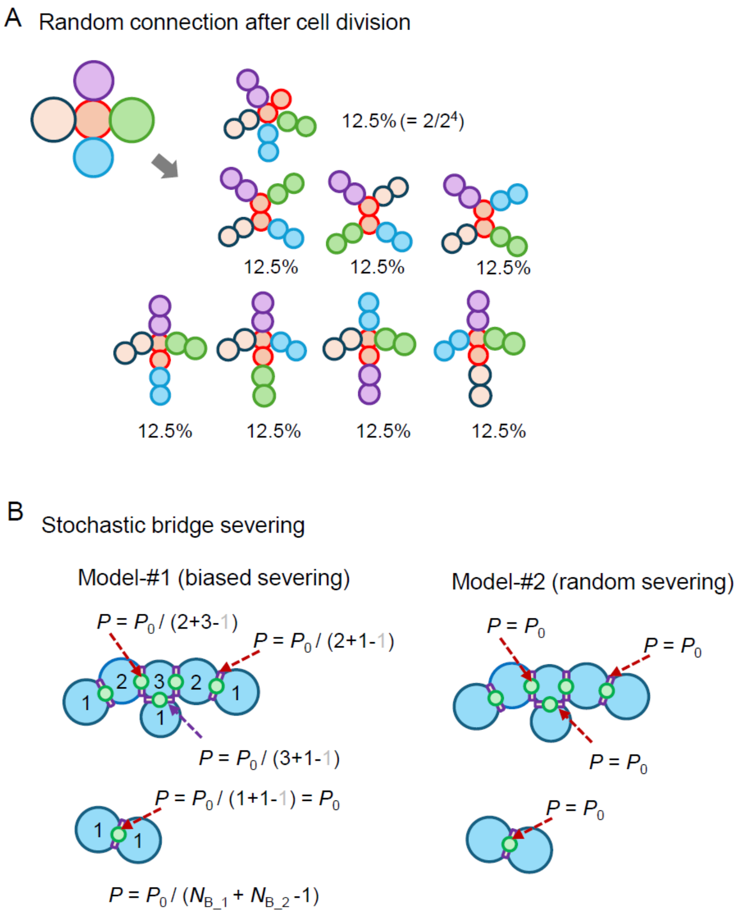


Figure S5 (related to Fig. 5): Definition of cell division and bridge severing in full model. A. The cell-cell connection after cell division is randomly determined. For instance, when a cyst comprises 5 cells where the centrale cell (red) has 4 bridges, 8 connection patterns can be stochastically generated. The two daughter cells originated from the same mother cell are shown in the same color. B. Two bridge severing scenarios are shown. In Model-#1, the bridge harbored by cells with more bridges is less severed. The number of the bridges for each cell is presented (e.g., 1, 2, 3). The severing probability for a certain bridge is determined by two terms: $P_0$ is a base probability, and the summation of the bridge number of the two cells connected by the bridge each other ($N_{B\_1}$ and $N_{B\_2}$). The precise definition of the probability is $P = P_0 / (N_{B\_1} + N_{B\_2} - 1)$, as exemplified in the panel. In Model-#2, the severing probability is constant ($P = P_0$) in a manner independent to the bridge number of the cells. This is equivalent to a "random severing" scenario.

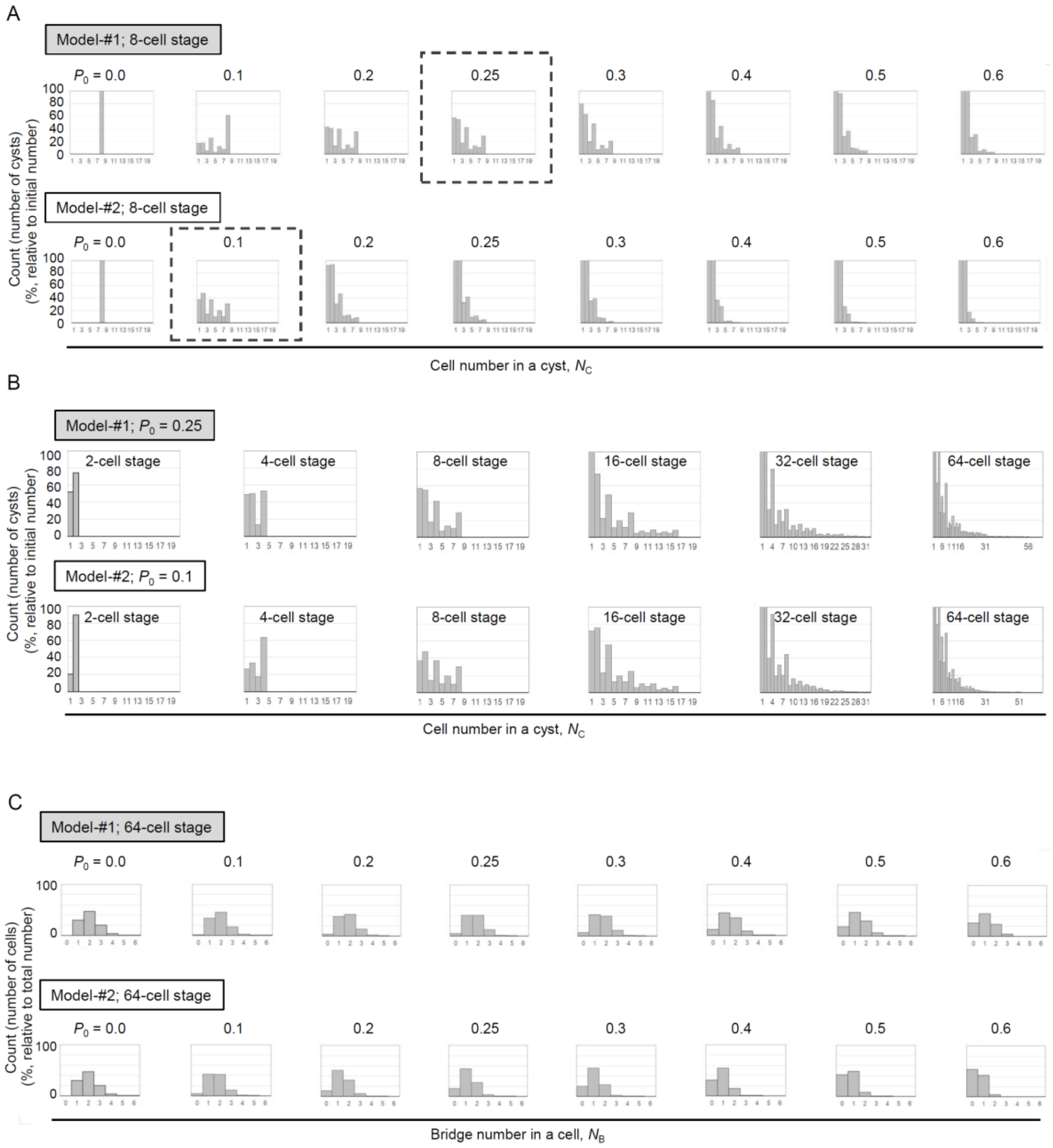


Figure S6 (related to Fig. 5): Severing probability recapitulating *in vivo* situation. A. The histograms of cell number in a cyst were generated in Model-#1 and -#2 which were defined in Fig. S5B. The simulations were performed under various values of the bridge severing probability ($P_0$ = 0.0 ~ 0.6), and the histograms at 8-cell stage are presented. The boxed histogram in Model-#1 corresponds to that in Fig. 5B, 8-cell stage. In comparison with our previous result (Ikami et al. 2023), the boxed histograms in the two models were most similar to the *in vivo* situation. B. The histograms of cell number in a cyst were generated in Model-#1 and -#2, according to the developmental stage (from 2-

cell stage to 64-cell stage). The bridge severing probabilities in the two models were adopted from the panel A ($P_0 = 0.25$ for Model-#1 and $P_0 = 0.1$ for Model-#2). C. The histograms of the bridge number in a cell were generated in Model-#1 and #2, at 64-cell stage. The bridge severing probability was varied ($P_0 = 0.0 \sim 0.6$).

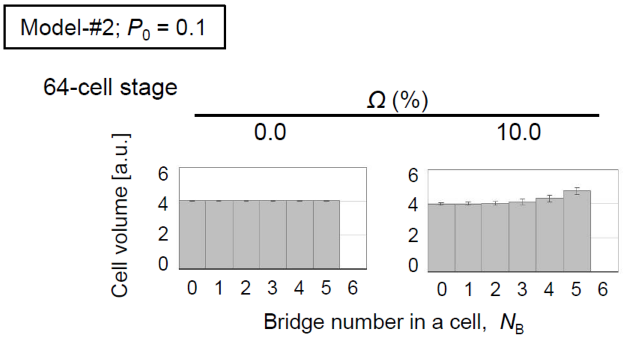


Figure S7 (related to Fig. 6): The histograms of the bridge number in a cell are presented in a manner similar to Fig. 6A-i, except that Model-#2 was adopted.
